# Switchable genome editing *via* genetic code expansion

**DOI:** 10.1101/349951

**Authors:** Toru Suzuki, Maki Asami, Sanjay G. Patel, Louis Y. P. Luk, Yu-Hsuan Tsai, Anthony C. F. Perry

## Abstract

Multiple applications of genome editing by CRISPR-Cas9 necessitate stringent regulation and Cas9 variants have accordingly been generated whose activity responds to small ligands, temperature or light. However, these approaches are often impracticable, for example in clinical therapeutic genome editing *in situ* or gene drives in which environmentally-compatible control is paramount. With this in mind, we have developed heritable Cas9-mediated mammalian genome editing that is acutely controlled by the cheap lysine derivative, Lys(Boc) (BOC). Genetic code expansion permitted non-physiological BOC incorporation such that Cas9 (Cas9^BOC^) was expressed in a full-length, active form in cultured somatic cells only after BOC exposure. Stringently BOC-dependent, heritable editing of transgenic and native genomic loci occurred when Cas9^BOC^ was expressed at the onset of mouse embryonic development from cRNA or *Cas9*^*BOC*^ transgenic females. The tightly controlled Cas9 editing system reported here promises to have broad applications and is a first step towards purposed, spatiotemporal gene drive regulation over large geographical ranges.

## Introduction

Many applications of genome editing by CRISPR-Cas9 (1–4) would benefit from being tightly controllable to minimize unintended, potentially harmful genome cleavage due to leaky nuclease activity. Such regulation would likely impact multiple contexts, including restricting genome editing to given anatomical sites in clinical somatic cell gene therapy (5,6) and gene drives (4,7). Gene drives duplicate a segment of genomic DNA *in vivo* independently of selection and in principle work in any sexually reproducing species so that all offspring inherit the gene drive segment (7). Given their speed and efficiency, gene drives have the potential to accelerate the dissemination of beneficial genetics in insects, crops and animals; they might streamline the introduction of homozygous mutations to study recessive alleles, eliminate destructive invasive species and agricultural pests, or to improve livestock rapidly and cheaply (for example, to prevent disease transmission), eliminating the need for protracted breeding programs (4). Recently, exceptionally high rates of gene drive transmission based on CRISPR-Cas9 genome editing were demonstrated in flying insect populations (8,9). However this remarkable efficiency mandates remarkable control.

Previous methods to control Cas9 activity (reviewed in Ref. 10) have involved modulating light (11–13) or temperature (13,14), but these approaches are difficult to regulate where barriers exist to light penetration (*eg* in larger organoids *in vitro* or environmentally) and are limited under conditions of temperature homeostasis (*eg* in mammals). Small molecule effectors including rapamycin, trimethoprim and (*Z*)-4-hydroxytamoxifen have also been used to control Cas9 activity in which, although each system differs, in general a domain is fused to Cas9 such that effector binding activates Cas9 through formation of functional protein (15–17), protein conformational change (18) or translocation into the nucleus (19).

However, these approaches often suffer from high background activities (16,18,19). For instance, a rapamycin-inducible system of gene activation caused a ~10-fold target increase in the absence of rapamycin (16) and the original demonstration of Cas9 switching by conformational change exhibited background editing of ~10%; refinement to reduce the background significantly reduced on-target editing (18). Moreover, rapamycin and trimethoprim are antibiotics with the potential to alter the microbiome, and as potential drivers of antimicrobial resistance, their deployment is likely to be problematic. (Z)-4-Hydroxytamoxifen, which is the major active metabolite of the anticancer agent, tamoxifen, is an antagonist of the estrogen receptor, hemolytic towards human erythrocytes, an endocrine disruptor and aquatic toxin (20–22). The cost of small molecule regulators in financial and environmental terms is also likely to be prohibitive on large scales.

Clearly, Cas9 regulation by other means will be advantageous or essential for many of its potential applications. To address this, we considered that a cheap amino acid such as the lysine derivative, H-Lys(Boc)-OH (BOC), might be harnessed for Cas9 control. BOC can be incorporated into proteins of interest by genetic code expansion (23), suggesting that a novel tier of regulating Cas9 might be achieved by generating a functional Cas9 variant whose expression (translation) and activity depended on the presence (incorporation during translation) of BOC. We therefore set out to evaluate the control of Cas9 by genetic code expansion in mammalian heritable genome alteration as a first step to the stringent control of gene drives and other applications.

## Results

### Establishing genetic code expansion in mouse embryos

As a starting point, we characterized genetic code expansion to control eGFP expression in transgenic mouse embryos. To this end, an orthogonal pyrrolysyl-tRNA synthetase/tRNA (PylRS/*Pyl* tRNA) pair was selected to direct incorporation of BOC into an amber stop codon at position 150 of *eGFP*, giving *eGFP*^*N150B*^ (24); we anticipated that in this system, translation of *eGFP*^*N150B*^ would be terminated prematurely (at codon 150) in the absence of BOC and functional eGFP^N150B^ protein produced only following BOC exposure (Figure 1A-D). Because we were interested in modulating protein activity in a mouse model, we designed an *eGFP*^*N150B*^ transgene to express each of the components required for ubiquitous, constitutive expression of the eGFP^N150B^ system (PylRS, *Pyl* tRNA and eGFP^N150B^; Supplementary Figure S1A) and coinjected it with wild-type (wt) mouse B6D2F1 sperm into B6D2F1 metaphase II (mII) oocytes (Supplementary Figure S1B). This method of transgenesis by intracytoplasmic sperm injection (ICSI) typically yields >80% of transgene-expressing preimplantation embryos after ~4 days of development (E3.5) (25,26). Following *eGFP*^*N150B*^ transgene injection, some of the resultant embryos (28.9%; *n*=44) clearly fluoresced after 4 days of culture in media supplemented with 10 mM BOC for the final 24 h, but only ~2% fluoresced (with a weak or partial signal; *n*=46) when BOC was omitted (*p*<0.001; Supplementary Figure S1C,D). This suggested that the orthogonal system could enable BOC-inducible eGFP^N150B^ expression in embryos. To enhance spatiotemporally regulated embryonic expression, we synthesized all RNA components of the BOC system (*ie PylRS* cRNA, *eGFP*^*N150B*^ cRNA and *Pyl* tRNA) *in vitro* and coinjected them into B6D2F1 mII oocytes followed by exposure to different concentrations of BOC (Supplementary Figure S2A). These injections were analogous to *eGFF*^*N1501B*^ transgenesis (Supplementary Figure S1A,B) except that RNA was delivered directly rather than relying on transcription of transgene DNA. All injected oocytes continually exposed to 1 mM BOC expressed readily-detectable green fluorescence (*ie* eGFP^N150B^) after 5 and 24 h (Supplementary Figure S2B-D). Thus, RNA injection resulted in efficient BOC-inducible eGFP^N150B^ expression in oocytes, paving the way for its application to other proteins.

**Figure 1.**
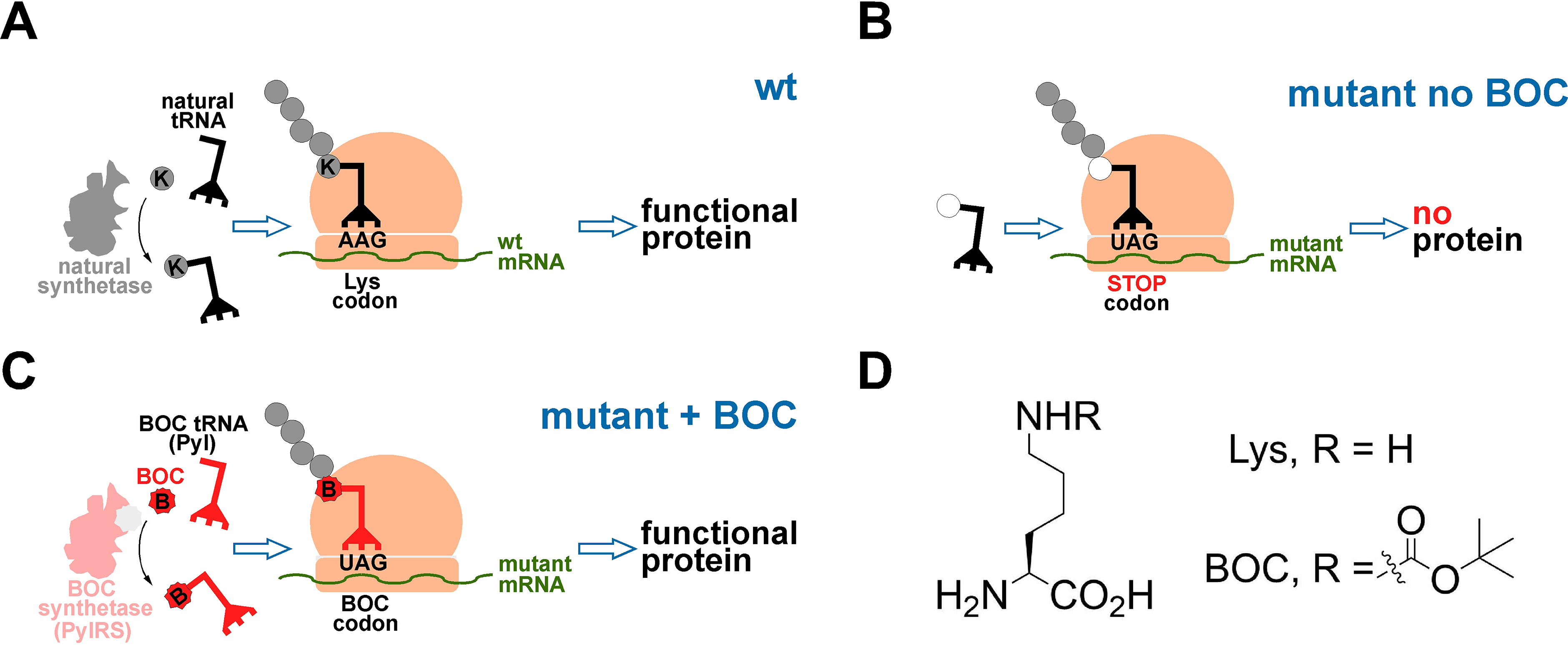
Schematic diagrams depicting natural translation and translation in the BOC system. (**A**) the natural incorporation of lysine (Lys, K) and (**B**) termination of translation at a stop codon. (**C**) When introduced ectopically, the orthogonal aminoacyl-tRNA synthetase, PylRS, can attach the non-physiological amino acid, BOC to its orthogonal tRNA, which decodes the stop codon, UAG, allowing BOC incorporation into a nascent polypeptide chain during translation. (**D**) Chemical structures of Lys and BOC.

### Genetic code expansion enables regulation of Cas9 activity *via* an orthogonal amino acid in tissue culture

We wished to evaluate whether genetic code expansion could be harnessed so that the RNA-guided nuclease activity of Cas9 (1,2,27) could be controlled by adding BOC. Since Cas9 residues can be substituted with a bulky Lys derivative without abolishing endonuclease activity (11), we tested Cas9 in which K510 or K742 were replaced with BOC residues. To evaluate this system, HEK293 cells were first transfected with constructs encoding *eGFP* gRNA, eGFP fused to a destabilization domain (eGFP-DD)(28), *Pyl* tRNA, PylRS and Cas9^K510B^, Cas9^K742B^ or Cas9^K510B,K742B^. Transfectants exposed to BOC expressed Cas9 mutants accompanied by a marked reduction of eGFP protein levels, suggestive of BOC-induced Cas9^K510B^ *eGFP* gene targeting activity (Figure 2A). Cas9^K510B^ activity was induced by BOC in the micromolar range in a dose-dependent manner (Figure 2B, *n*=3) and Cas9^K510B^ disappearance post-induction was accelerated when it was fused to the destabilization domain (Cas9^K510B^-DD in Figure 2C,D; *p*=0.002, *n*=3). We accordingly evaluated induction of Cas9 mutant activity by BOC in early embryos.

**Figure 2.**
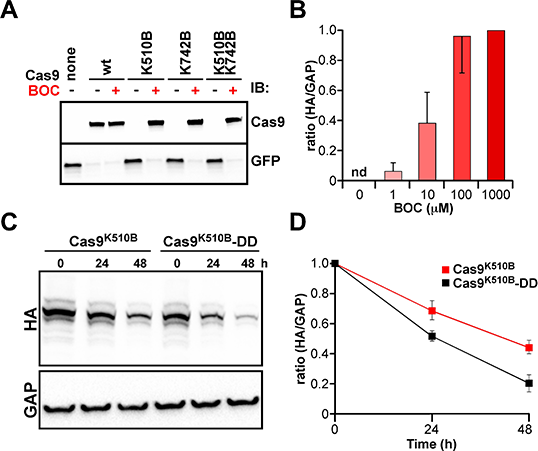
Regulation of Cas9 activity by BOC in mammalian *in vitro* cell culture. (**A**) Immunoblotting (IB) of HEK293 cells co-transfected with a construct expressing the HA-tagged Cas9 mutant indicated plus another construct encoding eGFP-DD, PylRS, gRNA and *Pyl* tRNA. Transfectants were incubated in the presence (+) or absence (−) of 1 mM BOC for 24 h before protein detection with antibodies against HA (Cas9) or GFP. Upper and lower panels represent cropped regions of the same representative (*n*=3) immunoblot, whose corresponding uncropped, full-length versions are presented in Supplementary Figure S3B. (**B**) Induction of Cas9^K510B^ following transient transfection of HEK293 cells and incubation for 24 h in the presence of the indicated concentrations of BOC. Induction was inferred by determining protein levels by immunoblotting with antibodies against HA (Cas9^K510B^) or GAPDH (GAP) and HA/GAP ratios normalized (value = 1.0) against the ratio at 1,000 μM BOC. nd, not detected. (**C**) Stability of Cas9 variants in HEK293 cells. Cells were transfected with constructs for PylRS, PylT, Cas9^K510B^ or Cas9^K510B^-DD and cultured in the presence of 1 mM BOC for 24 h. Transfectants were then washed with DMEM three times and analyzed after a further 0, 24 or 48 h by immunoblotting (IB) with antibodies against HA (for Cas9, which has an HA-tag) or a GADPH (GAP) loading/transfer control. Upper (full length) and lower (cropped) panels correspond to the same representative (*n*=3) immunoblot. (**D**) Plots of data from (**C**), in which the ratio of HA/GADPH signals at 0 h was set to 1.0 for each series. Error bars (s.d.) were from experimental triplicates. For the difference between Cas9^K510B^ or Cas9^K510B^-DD after 48 h, *p*=0.002 (*n*=3).

### BOC-switchable endogenous transgene editing in one-cell embryos

Oocytes were coinjected with *Pyl* tRNA and cRNAs encoding PylRS and Cas9^K510B^, Cas9^K742B^ or Cas9^K510B,K742B^, incubated for 4~5 h to allow cRNA-encoded protein expression (29) and then coinjected with *eGFP* gRNA plus sperm (30) from males homozygous for a ubiquitously-expressed single-copy *eGFP* transgene (31) previously produced by us (Figure 3A). Mutant Cas9 protein expression was enabled by exposure to BOC immediately after RNA injection for 4~5 or 24 h (Figure 3A). In these experiments (with Cas9^K510B^ unless stated otherwise), intracytoplasmic sperm injection (ICSI) was expected to produce embryos that fluoresced unless the *eGFP* gene had successfully been inactivated by editing (3). As anticipated, control embryos lacking one or more of the BOC system components all contained brightly fluorescent blastomeres because editing had not taken place and the paternal (sperm-derived) *eGFP* transgene was therefore expressed (Figure 3B,C). By contrast, when all BOC components were present and embryos exposed to 1 mM BOC, bright fluorescence was eliminated (Figure 3B,C). This clearly indicated that the *eGFP* gene had been inactivated with a high efficiency. Development to the blastocyst stage was reduced following 1 mM BOC exposure for 24 h but it was more efficient following briefer (4~5 h) BOC exposure (Figure 3D). *eGFP* mRNAs were intact in fluorescing blastocysts (*n*=12) but in all tested non-fluorescing blastocysts, sequencing revealed that transcripts corresponding to *eGFP* contained one or more mutations corresponding to the targeted region (*n*=8; Supplementary Figure S4A). Analogous experiments with cRNA encoding a different Cas9^BOC^ mutant, Cas9^K742B^, followed by 4~5 h BOC exposure also efficiently yielded non-fluorescing blastocysts (*n*=38; Figure 3E). Induced editing in the system also worked with the double mutant, Cas9^K510B,K742B^ (*n*=24), albeit less efficiently; 97.4% for Cas9^K742B^ (Figure 3E) *vs* 50.0% for the double mutant (Figure 3F). These data show that BOC system component levels compatible with embryo development induced efficient editing by Cas9^K510B^ and Cas9^K742B^. Furthermore, the developmental read-out clearly demonstrates that any truncated Cas9 protein produced in the absence of BOC, or pan-transcriptomic amber-codon read-through of endogenous mRNAs during BOC treatment, had limited cytotoxicity, because even though mouse preimplantation embryos are exquisitely sensitive to developmental perturbation, it did not prevent high rates of development. Moreover, editing was absolutely BOC-dependent, showing that in these experiments the system exhibited little, if any, leakiness.

**Figure 3.**
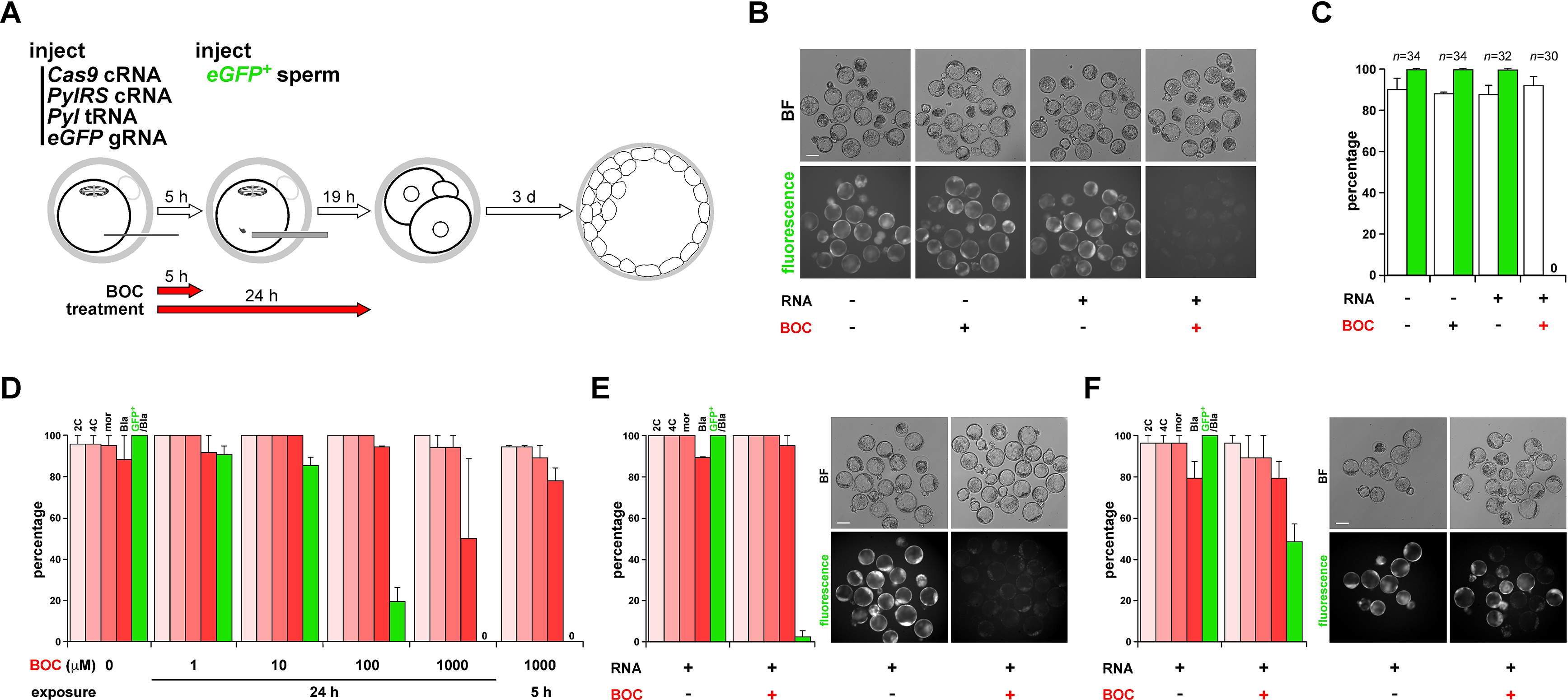
Genetic code expansion allows BOC-inducible, Cas9-mediated genome editing in mouse preimplantation embryos. (**A**) Schematic diagram depicting the experimental strategy: serial injection first of a mouse metaphase II (mII) oocyte with the indicated RNAs and 4-5 h later by an *eGFP*+ sperm carrying a ubiquitously-expressed *eGFP* transgene which fertilizes the oocyte. BOC (1 mM) is either omitted from the culture or included for the durations indicated after RNA injection. Oocytes were injected with a mixture of *Cas9*^*K510B*^ cRNA (600 ng/μl), *Pyl* tRNA (1,200 ng/μl) and *PylRS* cRNA (600 ng/μl) and 5 h later by *eGFP* gRNA (200 ng/μl) together with the *eGFP*+ sperm. (**B**) Vertically paired bright field (BF, upper) and fluorescence images of blastocysts produced by the method of (**A**) on the fourth day of culture (E4.0). Oocytes had been injected (+) with RNA, or RNA injection omitted, followed by exposure to BOC for 5 h (+) or not as indicated. Bar, 100 μm. (**C**) Histograms showing the percentage (±s.e.m.) of 1-cell embryos of (**B**) developing to the blastocyst stage at E4.0 (open columns) and the percentages of blastocysts that fluoresced green (green columns), with embryo numbers (*n*) given above columns. (**D**) Histograms showing the percentage (±s.e.m.) of 1-cell embryo development to 2-cell (2C), 4-cell (4C), morula (mor) and blastocyst (Bla) stages following serial injection first of mII oocytes with a mixture of *Cas9^K5I0B^* cRNA (600 ng/μl), *Pyl* tRNA (1,200 ng/μl) and *PylRS* cRNA (600 ng/μl) and 5 h later by *eGFP* gRNA (200 ng/μl) together with an *eGFP*+ sperm. Incubation was in media containing BOC at the concentrations indicated for 5 or 24 h after the first injection (*ie* respectively 5 h before ICSI and 19 h after). Percentages of blastocysts that were green fluorescent are represented by green columns. (**E**) Histograms (left) showing the percentage (±s.e.m.) of 1-cell embryos as per (**D**) except that *Cas9^K742B^* cRNA was injected instead of *Cas9^K5I0B^* cRNA and incubation was in the presence (+) or absence (−) of 1 mM BOC for 5 h after the first injection. Vertically paired images of E4.0 embryos are to the right, with culture in the presence (+) or absence (−) of 1 mM BOC for 5 h after the first injection. Bright field (BF) is shown (upper) with eGFP (fluorescence). Scale bar, 100 μm. (**F**) As per (**E**) except that cRNA encoding the *Cas9^K5I0B,K742B^* double mutant was injected instead of *Cas9^K742B^* cRNA.

### Heritable, BOC-dependent transgene editing

We next assessed editing heritability in the BOC-inducible system by performing a Cas9^K510B^ series in which *Pyl* tRNA- and cRNA-injected oocytes were immediately cultured in 1 mM BOC for 4~5 h, injected with *eGFP* gRNA plus *eGFP*+ sperm (Figure 3A) and the resulting 2-cell embryos non-selectively transferred to surrogate mothers. When injection of RNA system components and/or exposure to BOC were omitted, the resulting control offspring fluoresced green (Figure 4A,B), which was expected given that an intact *eGFP* allele was present in all fertilizing sperm. However, only 1/18 (5.6%) of pups derived from test embryos following BOC induction were fluorescent, suggestive of efficient editing (Figure 4A,B). Genomic sequencing of the targeted *eGFP* region revealed that green fluorescent offspring possessed intact alleles (*n*=2), whereas non-all fluorescing offspring harbored at least one of a range of mutations (*n*=14; Figure 4C). Although individuals typically contained a mixture of edited *eGFP* sequences consistent with mosaicism, we did not detect a mixture of fluorescence and non-fluorescence in offspring, suggesting that editing had occurred efficiently at the 1-cell stage. These data suggest that BOC had efficiently induced, and was a stringent requirement for, Cas9^K510B^ genome editing activity and that the extent of editing in preimplantation embryos provided an accurate meter of editing in offspring.

**Figure 4.**
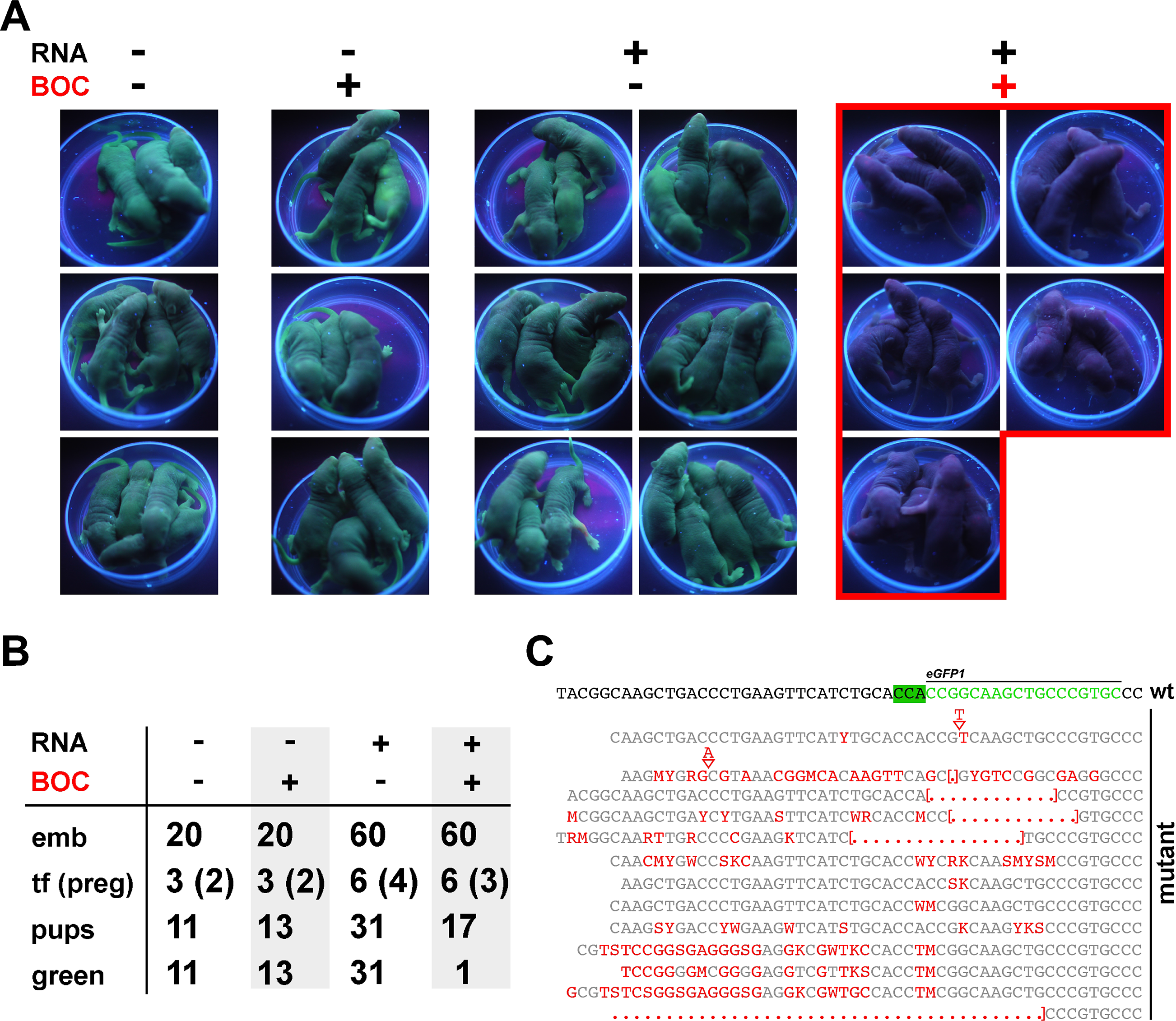
Heritable, BOC-inducible editing of an endogenous *eGFP* transgene by cRNA-encoded Cas9^K510B^. (**A**) Offspring produced by serial injection of C57BL/6 (B6) mII oocytes with RNA (+) or not injected (−), and incubated in the presence (+) or absence (−) of 1 mM BOC for 4-5 h as indicated, followed by injection of an *eGFP+* sperm. Two-cell embryos were transferred non-selectively to recipients and the resulting offspring imaged on postnatal day 3 under UV illumination. Injected RNA (RNA) comprised a mixture of *PylRS* cRNA (600 ng/μl), *Cas9^K5I0B^* cRNA (600 ng/μl), *Pyl* tRNA (1,200 ng/μl) and *eGFP* gRNA (200 ng/μl). (**B**) Developmental efficiency and green fluorescence in offspring following BOC-inducible editing by Cas9^K510B^ of (**A**) in the presence (+) or absence of 1 mM BOC. The number of embryos (embryos) transferred to recipients (tf) is given relative to the number of recipients that became pregnant (preg). *Died perinatally. (**C**) Sequences of genomic PCR products encompassing the targeted region of *eGFP*, derived from nonfluorescing blastocysts in the experiment of (**A**). The sequence corresponding to the *eGFP* target (green) plus adjacent sequences are displayed on the top row and mutants beneath, with the proto-spacer adjacent motif (PAM) highlighted in green. Mutations are indicated in red type-face. All (*n*=12) non-fluorescing blastocysts examined had undergone mutagenesis whereas no controls (*n*−12) in which either RNA or BOC had been omitted harbored mutations.

### BOC-dependent heritable editing of native alleles

We next determined whether, in an analogous protocol, BOC could induce Cas9^K510B^-mediated editing of native alleles of the mouse strain, B6. To this end, we selected genes encoding the testis-determining factor, Sry (32), or tyrosinase (Tyr), which confers dark fur pigmentation in B6 mice (33). Targeting *Sry* with control Cas9 (lacking a BOC codon) and a mixture of two gRNAs (Supplementary Table S1) yielded 16 females of which five (31.3%) carried *Sry* harboring a mutation, implying that *Sry* inactivation had resulted in sex reversal. When we employed Cas9^K510B^ in the absence of BOC, 0/7 of the resulting females carried *Sry*, but when BOC was included, 2/10 (20.0%) females were *Sry*-positive (Figure 5A) and in both cases *Sry* had undergone frame-shifting edits (Supplementary Figure S4B). Editing the coat-color determinant, *Tyr* by control Cas9 and three gRNAs (Supplementary Table S1) had previously resulted in 92.9% (*n*=14) of offspring with altered coat-color phenotypes (34). A similar efficiency (93.3%; *n*=15) of *Tyr* editing was achieved by BOC-dependent Cas9^K510B^-mediated targeting (Figure 5B,C); sequence analysis of coat-color mutants (*n*=6) revealed that all contained one or more frame-shifting *Tyr* gene indels in the targeted region (Supplementary Figure S4C) with a pattern of sequence mosaicism analogous to those observed for *eGFP* editing (Supplementary Figure S4A). This suggests that BOC-induced editing of endogenous loci by Cas9^K510B^ occurred at comparably high efficiencies to editing mediated by control Cas9.

**Figure 5.**
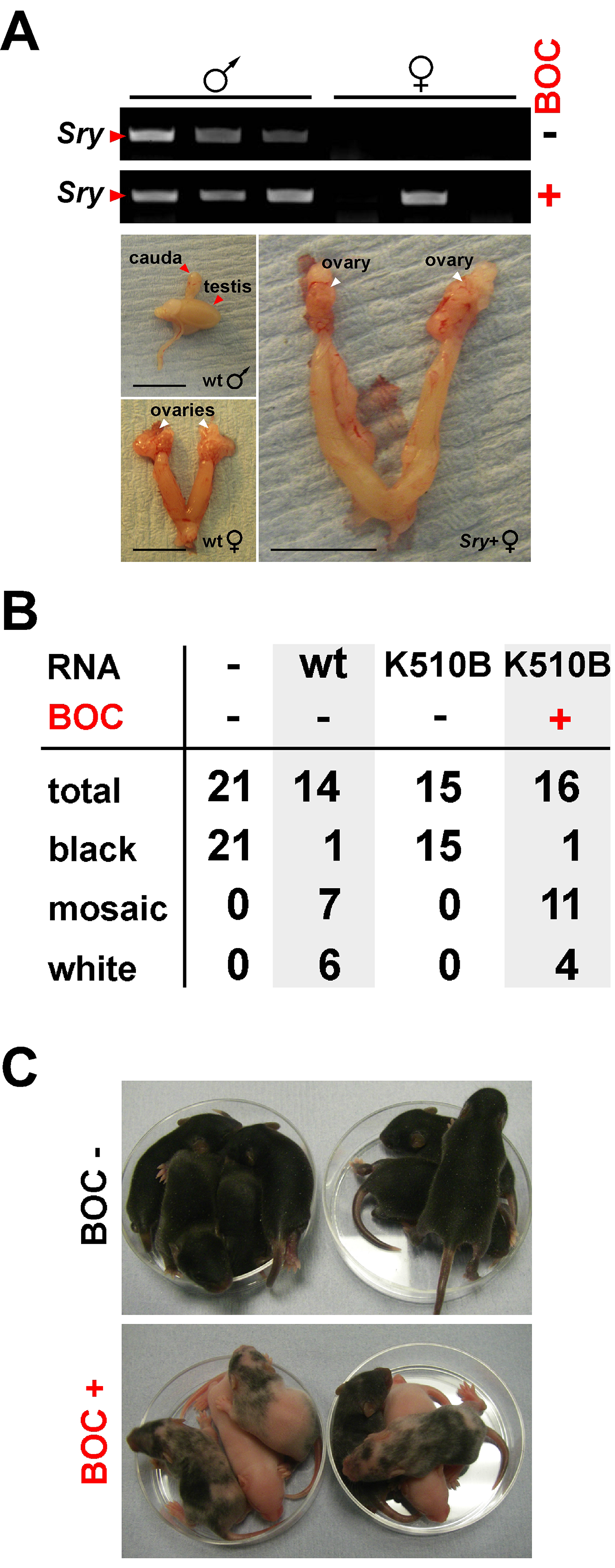
Heritable, BOC-induced editing of native genomic alleles by Cas9^K510B^ expressed from cRNA. (**A**) BOC-induced editing of the sex-determining gene, *Sry* following injection of wt C57BL/6 (B6) mII oocytes with B6 sperm and a mixture of *PylRS* cRNA (600 ng/μl), *Cas9^K5I0B^* cRNA (3,300 ng/μl), *Pyl* tRNA (1,200 ng/μl) and three gRNAs (800 ng/μl) (Supplementary Table S1). Results of ear-punch genomic PCR for *Sry* are presented (upper) for offspring generated when BOC had been included (+) or omitted. Internal reproductive organs are shown for wild type (wt) and sex-reversed adults. Scale bar, 5 mm. (**B**) Developmental efficiency and phenotypes of offspring produced by ICSI with wt B6 sperm and oocytes as per (**A**), except that the mixture of three gRNAs targeted *Tyr* (Supplementary Table S1). The total number of offspring (total) is given with numbers exhibiting different coat-color phenotypes. Data reflect at least two experiments per target. (**C**) Offspring of (**B**) at 7 d.

### Inducible genome editing *via* Cas9^BOC^ supplied by a mouse transgenic line

We extended this principle to evaluate BOC-induced editing by Cas9^K510B^ driven in transgenic mice by the *ZP3* promoter, *pZP3*, which is uniquely expressed in growing oocytes and has become a classical driver of transgene expression in the maternal germline (35,36). Comparison of expression between *pZP3-Cas9* and *pZP3-Cas9*^*K510B*^ transgenic lines is confounded by the differing sites of transgene integration in each, but within each line we noted a marked decline in expression in hemizygotes (containing a single transgene allele) from germinal vesicle stage oocytes, such that *pZP3* transgene mRNA was not detectable in mature, fertilizable mII oocytes of the type used in ICSI (Figure 6A). Thus, the levels of transgene-encoded *Cas9* and *Cas9*^*K510B*^ transcripts were extremely low.

**Figure 6.**
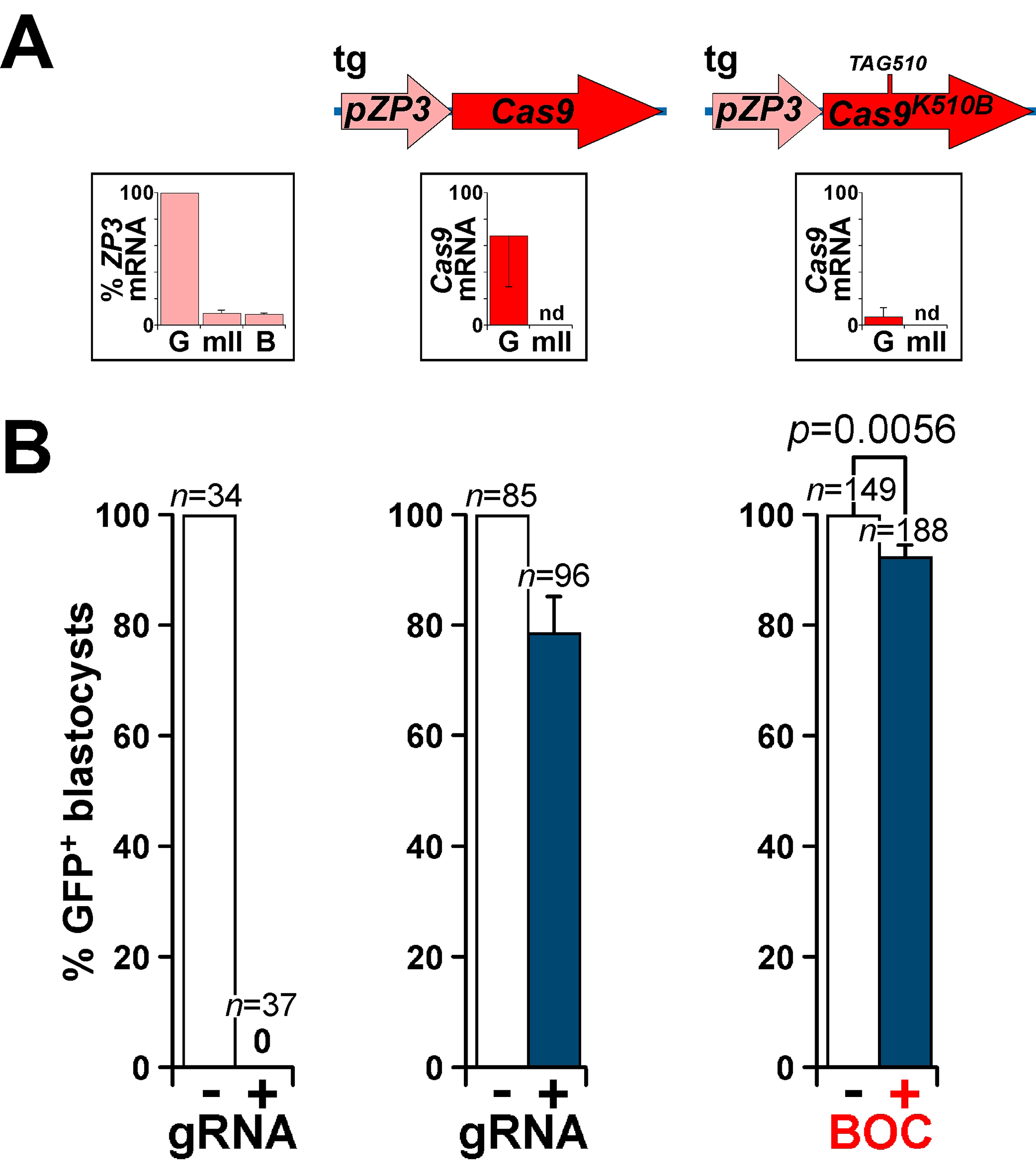
Heritable, BOC-induced editing of an endogenous paternal genomic *eGFP* transgene by endogenous maternal transgene-expressed Cas9^K510B^. (**A**) Structures of maternally-derived endogenous transgene (tg) encoding Cas9^K510B^ (right) or control Cas9, showing their corresponding mRNA expression levels ±s.e.m. (determined by qPCR; *n*=3) relative to levels of *H3f3* mRNA in immature germinal vesicle (G) and mature metaphase II (mII) oocytes. The activity relative to that in germinal vesicle oocytes (set at 100%) of the native *pZP3* promoter is indicated by similarly determining *ZP3* transcript levels (*n*=4) in wt oocytes and embryos (leftmost; B, E4.0 blastocyst). (**B**) Editing of a paternally-derived transgenic *eGFP* allele following *eGFP+* sperm injection (ICSI) as determined by the percentage of blastocysts (±s.e.m.) that fluoresced on E4.5. In controls, wt C57BL/6 (B6) mII oocytes (leftmost) or mII oocytes from *pZP3-Cas9* transgenic females [centre, corresponding to the tg structure of (**A**), above] were injected with *eGFP+* sperm and a mixture of *Cas9* cRNA (3,300 ng/μl) and *eGFP* gRNA (800 ng/μl). Also (rightmost), mII oocytes from *pZP3-Cas9^K5I0B^* transgenic females (tg structure above) were injected with *eGFP+* sperm and a mixture of *PylRS* cRNA (600 ng/μl), *Pyl* tRNA (1,200 ng/μl) and *eGFP* gRNA (800 ng/μl) (Supplementary Table S1) but no cRNA encoding Cas9^K510B^. Incubation either omitted BOC (−) or included 1 mM BOC for 5 h (+).

When mII oocytes from *pZP3-Cas9* hemizygous transgenic females were injected with *eGFP*+ sperm plus gRNA targeting *eGFP* (but no *Cas9* cRNA), 21.2±6.9% of the resulting blastocysts (*n*=96) underwent editing as judged by loss of fluorescence (Figure 6B); editing was confirmed by sequencing (Supplementary Figure S4D). Genome editing had therefore been mediated by endogenous transgene-encoded Cas9 in mII oocytes from hemizygous transgenic females.

We next evaluated whether this might be extended to a BOC-inducible system by injecting mII oocytes from*pZP3-Cas9*^*K510B*^ hemizygous transgenic females with *Pyl* tRNA and cRNA encoding PylRS (but without cRNA encoding Cas9^K510B^), followed by *eGFP*+ sperm plus gRNA targeting *eGFP*. Blastocyst development rates were similar in both groups: 63.3±3.1% of injected oocytes (*n*=235) developed in the absence of BOC and 64.4±3.8% (*n*=281) with it (*p*=0.815). When BOC was included, 7.4±2.3% of the resulting blastocysts (*n*=188) lacked fluorescence (including one mosaic) and contained edited *eGFP* transcripts, whereas all (*n*=149) of the blastocysts fluoresced brightly when it was omitted (*p*=0.006; Figure 6B). Given that the control editing rate was 21.2% (see above), BOC-dependent editing by *Cas9^K510B^* transgene-expressing females had therefore occurred at ~34.9% of the control rate (7.4% for *Cas9^K510B^ vs* 21.2% for the control; Figure 6B).

## Discussion

This work demonstrates controllable, heritable genome editing in mouse embryos by inducing Cas9 activity with the cheap, non-physiological (in mammals) amino acid, Lys(Boc) (BOC) *via* genetic code expansion. Genetic code expansion has previously been employed for the incorporation of non-physiological amino acids other than BOC, always into the bystander reporter protein, GFP *in vivo* (37–39). We here demonstrate that the BOC system applied to Cas9 is compatible with efficient full-term development even when it is active at an early and exquisitely sensitive phase of embryogenesis. This not only provides a developmental indication of limited or surmountable toxicity due to broad genomic amber codon read-through (at least when it occurs during early mammalian embryogenesis) but argues against extensive off-target editing in the system (40). In contrast to the higher background activities or restricted applicability of many switchable systems (11–13,16,18,19), this work accordingly demonstrates low or zero Cas9^BOC^ activity in the absence of BOC; without BOC, Cas9^BOC^ activity was undetectable under the conditions of these experiments, suggesting that the system was subject to little, if any, leakiness.

The BOC-inducible principle should also apply to modified Cas9 activities such as ‘nuclease-dead’ Cas9 (dCas9) derivatives fused to transcriptional modifiers (DNA methyltransferase, histone acetyltransferase, transcription factor activation domains, *etc*.) to enable reversible, dynamic transcriptional gene control and epigenetic editing (modification) on physiological time-scales (within minutes) (41–43). BOC would have clear potential advantages over other small molecule effectors, which may either interfere with the fused epigenetic modifier domain or be occluded by it, necessitating case-by-case validation. Moreover, some effectors (*eg* tamoxifen) inherently contribute to chromatin remodeling, complicating data interpretation (44,45).

BOC-inducible Cas9 in gametes or embryos near the time of fertilization also promises to mediate tight gene drive regulation. Although potentially powerful, gene drives may have unforeseen and uncontrollable consequences (4). The Cas9 system described here addresses this, and because BOC is a cheap amino acid, it could potentially be used environmentally to control gene drive expression. We generally used 1 mM BOC, but even in a non-optimized *in vitro* system, a 1,000-fold lower BOC concentration worked detectably (Figure 2B). Lower concentrations are expected to exert fewer co-lateral effects and reduce costs. Thus, BOC (or something analogous) promises to enable gene-drive activity that can be controlled over large geographic ranges, in principle restricting them to areas of BOC exposure. Gene-drive switchability will also reduce the emergence of resistance by periodically removing the selectable advantage of gene drive neutralization; once the gene drive is inactive, it cannot be selected against. There is no reason *a priori* why this approach should be limited to editing by Cas9, and other nuclease frameworks (*eg* Cpf1) should be amenable to analogous regulation.

In addition, genome editing is non-trivial in large animals, whose importance in the provision of biomedical models and clinical reagents (*eg* for xenotransplantation) is increasing (46). By streamlining safe target locus homogenotization (*ie* allelic replacement to homozygosity) in founders generated by nuclear transfer cloning, pronucleus or sperm injection or *via* ES/iPS cells, gene drive cassettes could reduce breeding programs and add flexibility, particularly impacting species such as pigs, where ES or naïve iPS cells have not been described. Gene drives could be activated by BOC in early embryogenesis following natural mating, obviating the need for microsurgery. The principle of Cas9 regulation by BOC should also apply to anti-Cas9 proteins (47), thereby producing a neutralizing effect on Cas9 activity that is itself controllable.

Moreover, BOC-regulated Cas9 (variants) may directly impact human biomedical applications including therapeutics (6); they would, for example, enhance spatiotemporal control of targeted viral integration and/or activity in somatic cell therapies (by restricting BOC to a given anatomical site and time) and mitigate against neutralization by preexisting anti-Cas9 antibodies (48). This control principle holds for proteins other than Cas9.

In sum, by extending orthogonal post-transcriptional regulation to functional, non-reporter genes, this work achieves a new tier of Cas9 control with the clear potential for broad utility in stem cell biology and suggests applicability to the control of diverse biological contexts *in vitro* and *in vivo*, to plants, insects and mammals, environmentally and in clinical practice.

## Materials and Methods

### Plasmid construction

Plasmid pEF1α-FLAG-PylRS-IRES-Neo-4xU6-PylT^U25C^ (a kind gift from Dr. Jason Chin) was used for the expression of pyrrolysyl-tRNA synthetase and pyrrolysyl tRNA_CUA_ and to direct the incorporation of BOC. Plasmid pCAG-eGFP has been described previously (25). Plasmids p3s-Cas9HC (Addgene plasmid #43945) and pU6-gRNA-GFP(5-10) were generous gifts respectively from Dr. Jin-Soo Kim and Prof. Anton Wutz; pU6-gRNA-GFP(5-10) expresses a gRNA targeting *eGFP* corresponding to the region between codons 5 and 10. We generated pCAG-eGFP^N150B^ using pCAG-eGFP as template in a PCR reaction with primers F1 and R1 (primer sequences are given in Table S1) to replace the N150 codon of *eGFP* with TAG (a B codon), as verified by sequencing with primers S1 and S2. To construct pCAG-eGFP-DD (encoding eGFP^T308D,S473D^), a 906 base pair (bp) fragment containing *eGFP-DD* (GeneArt, Regensburg, Germany) was introduced by Gibson assembly into *Eco*RI-digested pCAG-eGFP and verified by sequencing with primers S1 and S2. To generate pEF1α-FLAG-PylRS-CAG-eGFP(150TAG)-4xU6-PylT^U25C^ (designated SP44), the 8,286 bp *Bam*HI-*Sal*GI fragment of pEF1α-FLAG-PylRS-IRES-Neo-4xU6-PylT^U25C^ was joined by Gibson assembly to a 2,748 bp CAG-eGFP^N150B^ PCR amplimer generated with primers F4 and R4 from pCAG-eGFP^N150B^ and confirmed by sequencing with primers S1, S2 and S4. To construct plasmid p3s-Cas9HC^K510B^ (SP81), Gibson assembly was used to join the 5,862 bp *Kpn*I-*Bsr*GI fragment from p3s-Cas9HC to a PCR amplimer from p3s-Cas9HC containing Cas9(6-528) in which the codon for K510 had been replaced by TAG (B) using primers F5 and R5. The sequence of p3s-Cas9HC^K510B^ was verified by sequencing with primers S5, S6 and S7. To construct plasmid p3s-Cas9HC^K742B^ (SP82), the 6152 bp *Pml*I-*Bsr*GI fragment from p3s-Cas9HC was joined by Gibson assembly to a PCR amplimer from p3s-Cas9HC containing Cas9(514-745) (SP116) in which the codon for K742 had been replaced by TAG (B) using primers F6 and R6. The sequence of p3s-Cas9HC^K510B^ was verified by sequencing with primers S7, S8 and S9. Construct pU6-gRNA-GFP(51-56) was prepared from pU6-gRNA-GFP(5-10) and its sequence confirmed with primers S16 and S17. Plasmid pEF1α-FLAG-PylRS-CAG-eGFP-DD-4xU6-PylT^U25C^-U6-gRNA-GFP(5-10)-U6-gRNA-GFP(51-56) was constructed in four steps. First, Gibson assembly of the 8,286 bp pEF1α-FLAG-PylRS-IRES-Neo-4xU6-PylT^U25C^ *Bam*HI-*Sal*GI fragment to the 683 bp fragment (containing *IRES)* amplified with primers F2 and R2 from pEF1α-FLAG-PylRS-IRES-Neo-4xU6-PylT^U25C^, and the 758 bp fragment (containing *eGFP*) amplified with primers F3 and R3 from pCAG-eGFP generated pEF1α-FLAG-PylRS-IRES-eGFP-4xU6-PylT^U25C^; this was confirmed by sequencing with primers S1, S2 and S4. Secondly, Gibson assembly of the pEF1α-FLAG-PylRS-IRES-eGFP-4xU6-PylT^U25C^ *Bgl*II fragment with the 412 bp PCR product generated with primers F13 and R13 from pU6-gRNA-GFP(5-10) yielded pEF1α-FLAG-PylRS-IRES-eGFP-4xU6-PylT^U25C^-U6-gRNA-GFP(5-10), as confirmed by sequencing with primer S18. In the third step, Gibson assembly of the pEF1α-FLAG-PylRS-IRES-eGFP-4xU6-PylT^U25C^-U6-gRNA-GFP(5-10) *Bgl*II fragment with the 412 bp PCR product generated with primers F13 and R13 from pU6-gRNA-GFP(51-56) yielded pEF1α-FLAG-PylRS-IRES-eGFP-4xU6-PylT^U25C^-U6-gRNA-GFP(5-10)-U6-gRNA-GFP(51-56), which was confirmed by sequencing with primer S18. Finally, Gibson assembly of the 9,020 bp *Bam*HI-*Sal*GI fragment of pEF1α-FLAG-PylRS-IRES-eGFP-4xU6-PylT^U25C^-U6-gRNA-GFP(5-10)-U6-gRNA-GFP(51-56) and the 2,875 bp PCR amplimer generated using primers F4 and R4 from pCAG-eGFP-DD produced pEF1α-FLAG-PylRS-CAG-eGFP-DD-4xU6-PylT^U25C^-U6-gRNA-GFP(5-10)-U6-gRNA-GFP(51-56), as confirmed by sequencing with primers S1, S2 and S4. Plasmids were linearized prior to oocyte injection. In brief, the *PylRS* cassette was excised and inserted into the pCIneo vector (Promega) at its *NheI/EcoRI* site and *PylRS* cRNA synthesized *in vitro* using a T7 transcript kit (Invitrogen). The Pyl tRNA insert was subcloned as a 4x (U6-PylT) cassette into the vector, to give T7 promoter-directed expression. Plasmids were expanded and purified using EndoFree Plasmid Maxi Kit (QIAGEN) and digested with *Sfi*I/*Sap*I*Sal*lHF (SP44) or with *Xho*I for Cas9HC^K510B^ (SP81), Cas9HC^K742B^ (SP82) and Cas9HC^K510B,K742B^ (SP116). Plasmid p3S-Cas9HC^K510B^-DD, encoding Cas9HC^K510B^ (Cas9^K510B^ containing a C-terminal destabilization domain fusion) was constructed by Gibson assembly of the 7,320 bp p3S-Cas9HC^K510B^ *Xho*I-*Kpn*I fragment to a 233 bp fragment containing the destabilization domain (GeneArt). Fragments were gel-purified (Promega) and each was injected at 1.0 ng/μl into mII oocytes with sperm (25).

### Cas9 editing assay in HEK293 cells

HEK293 cells were grown at 37 °C in a 5% (v/v) CO_2_ atmosphere in DMEM plus GlutaMAX medium (Gibco) supplemented with 10% (v/v) fetal bovine serum for 24 h before transfection. Cells (2 × 10^5^/well in a 24-well plate) were transiently transfected with the appropriate p3s-Cas9HC variant and pEF1α-FLAG-PylRS-CAG-eGFP-DD-4xU6-PylT^U25C^-U6-gRNA-GFP(5-10)-U6-gRNA-GFP(51-56) using Lipofectamine 2000 (Life Technologies) according to the recommendations of the manufacturer and plated in parallel in media either further supplemented with 1 mM BOC or lacking BOC, or at the BOC concentrations indicated for Figure 2. After 24 h, cells were washed with 0.5 ml phosphate-buffered saline (PBS) and lysed with RIP A buffer (Sigma, R0278; 50 μl per well) containing protease inhibitor (Sigma, P8340) at 4 °C for 10 min. Lysates were pelleted (20,000 *g*, 10 min, 4 °C) and the supernatant (45 μl) added to 15 μl LDS sample buffer (Life Technologies). Samples were heated (95 °C, 10 min) and loaded onto a 4-20% Mini-PROTEAN TGX Stain-Free Protein Gel (Bio-Rad) for electrophoresis. The gel was imaged on a ChemiDoc XRS+ (Bio-Rad) and transferred to a nitrocellulose membrane using a Trans-Blot Turbo Transfer System (Bio-Rad). The membrane was stained with Ponceau S to confirm protein transfer, blocked with PBST (0.05% [v/v] Tween 20 in PBS) containing 5% (w/v) milk powder at 20 °C for 1 h, and then incubated (4 °C, overnight) with a primary mouse anti-GFP antibody (Thermo Fisher, #MA1-952, 1:500 [v/v] dilution). The membrane was then washed three times (10 ml PBST, 5 min per wash). All subsequent washing steps used this procedure. The membrane was then incubated in secondary anti-mouse antibody (Thermo Fisher, #32430, 1:1,000 [v/v] dilution) for 1 h at 20 °C, 1 h and then washed. The signal was developed by addition of Clarity Max™ Western ECL Substrate (Bio-Rad, #1705062). After imaging on a BioRad ChemiDoc XRS + system, the membrane was washed and incubated (~20 °C, 30 min) with Restore Western Blot Stripping Buffer (Thermo Fisher, #21059) and then washed, incubated (4 °C, overnight) with a primary mouse anti-HA antibody (Thermo Fisher, #26183, 1:2000 [v/v] dilution) before being incubated with the secondary antibody and processed for imaging as described above.

### Target gRNA synthesis

Target gRNAs were produced as previously described (49). Briefly, T7-gRNA-scaffold products were amplified by PCR using Pfx polymerase (Invitrogen) with Scaffold-Fwd and Scaffold-Rev primers and the T7 gRNA vector as template with PCR reaction parameters: 94 °C, 2 min followed by 25 cycles of 94 °C, 15 sec; 55 °C, 30 sec; 68 °C, 20 sec; final extension at 68 °C, 7 min. gRNA scaffold PCR product were gel purified (Promega) and used as a template using T7-target oligo and the amplified gRNA scaffold. gRNA was synthesized by PCR with Pfx using T7-target oligo primers (Supplementary Table S1), T7-fwd primer and scaffold-Rev primer with the parameters: 94 °C, 5 min followed by 25 cycles of 94 °C, 15 sec; 55 °C, 30 sec; 68 °C, 20 sec; final extension at 68 °C, 7 min. The amplification of single 130 bp PCR products was each case confirmed by 2% (w/v) agarose gel electrophoresis and products purified Promega) for following synthesis of gRNA *in vitro* (3) using the MEGAshortscript T7 Transcription Kit (Invitrogen) as recommended by the manufacturer.

### Collection and culture of oocytes and embryos

Animal procedures complied with the Animals (Scientific Procedures) Act, 1986 and experimental protocols were approved by the University of Bath Ethical Review Board. Wild-type mouse (*Mus musculus*) strains were bred from C57BL/6J (B6) and DBA/2 stocks in-house or supplied by Charles River (L’Arbresle, France). A 129SvJ line containing a single-copy *eGFP* transgene under the control of the *pCAG* promoter-enhancer has been reported previously (31). Oocytes were collected from 8-to 12-week-old females that had been super-ovulated by standard serial intraperitoneal injection of 5 IU pregnant mare serum gonadotropin (PMSG) followed 48 h later by 5 IU human chorionic gonadotropin (hCG) and held under mineral oil in humidified 5% CO_2_ (v/v air) at 37 °C, until required (29,31). Embryo culture *in vitro* was in KSOM under mineral oil in humidified 5% CO_2_ (v/v in air) at 37 °C.

### Sperm preparation and microinjection

Cauda epididymidal sperm from 8- to 12-week-old males were prepared as described previously (3,30,31). Sperm were resuspended in ice-cold nuclear isolation medium (NIM; ~0.5 ml per epididymis) and stored on ice or at 4 °C. Immediately prior to injection, ~50 μl of freshly-prepared sperm suspension was mixed 20 μl of polyvinylpyrrolidone (PVP, average *M*_r_ ≈ 360,000; Sigma-Aldrich) solution (15% [w/v]). DNA or gRNA solutions were immediately mixed with the sperm/PVP suspension to give each at a final concentration of 1 ng/μl and 200 ng/μl respectively in the suspension. For endogenous gene editing of *Tyr* and *Sry* loci in the BOC system, the gRNA working concentration was 800 ng/μl. Approximately 2.0~2.5 μl of suspension containing a single sperm head was injected (ICSI) within ~60 min of DNA/gRNA mixing into mII oocytes held in droplets of M2 under mineral oil, essentially as described (25,30,31). Where indicated, mII oocytes that had been injected with RNA were injected with sperm heads (ICSI) 3~4 h later.

### Synthesis of RNA. *in vitro* and microinjection into mII oocytes

The preparation of 5′-capped and polyadenylated in vitro transcripts was from cRNA synthesized from linearized plasmid template DNA in a T7 mScript™ Standard mRNA Production System (Cellscript, USA) as previously described (29,31). Synthesis *in vitro* of gRNA and tRNA was with the MEGAshortscript T7 RNA synthesis kit (Invitrogen) according to the instructions of the vendor. Each took place at 37 °C for 3~4 h in a 20 μl total reaction volume containing DNA template and T7 RNA polymerase. RNAs were precipitated, dissolved in nuclease-free water, quantified on a Nanophotometer and stored in aliquots at ^-^ 80 °C until required. RNA solutions were diluted as appropriate with sterile water and injected within 1 h of thawing via a piezo-actuated micropipette into mII oocytes, or in the case of gRNA, in the presence of sperm. Working concentrations of RNA injected into mII oocytes were: 600 ng/μl for gRNA, 100 ng/μl for cRNA encoding non-mutant Cas9, 600 ng/μl for cRNA encoding PylRS, eGFP^N150B^ or mutant Cas9, and 1,200 ng/μl for *PylT^U25C^* tRNA (*Pyl* tRNA). For BOC-induced editing of endogenous genomic *Sry* and *Tyr* sequences, the working concentrations of cRNA encoding mutants was 3,300 ng/μl and for gRNA, 800 ng/μl. We estimate that 5~10 pl were typically injected per mII oocyte.

### Reverse transcriptase PCR

Total RNA from embryos and tissues was extracted using the TRI reagent (Sigma) according to the instructions provided. From each total RNA sample, 800 ng were used for first strand cDNA synthesis by Superscript III Reverse transcriptase (Invitrogen). Following synthesis, cDNA was treated with RNase H (Invitrogen) for 20 min at 37 °C and 1 μl of resultant cDNA used to charge PCR reactions with the appropriate target primers (Supplementary Table S1).

### Embryo transfer

Embryonic day (E1.5) 2-cell embryos were transferred to the oviducts of pseudo-pregnant CD-1 females at day 0.5 (*ie* plugged females that had been mated with vasectomized males the previous night). Where appropriate, pups were delivered by Cesarian section and fostered by CD-1 females.

### Genotyping

Genotyping was performed on ear-punch biopsies collected at weaning as described previously (3) and digested at 55 °C for 5 h in 100 μl of a lysis buffer containing 10% (w/v) sodium dodecyl sulfate and 2 mg/ml proteinase K (Sigma). One microliter of a 1:10 (v/v) dilution of each genomic DNA sample was used for genotyping by PCR in a 10 μl reaction volume. For single blastocyst genomic PCR, individual blastocysts (E4.5-E5.5) were each collected into a PCR tube containing 0.5% (w/v) sarcosyl and flash-frozen in liquid nitrogen before being subjected to target PCR amplification with primer sequences as given in Supplementary Table S1. Where appropriate, PCR products were gel-purified for sequence analysis.

### Fluorescence imaging

Following oocyte injection, fluorescence of embryos was visualized on an IX71 (Olympus, Japan) microscope equipped with an AndroZyla cMOS camera and OptoLEP illumination system (Cairn Research, UK) as previously described (31). Images were processed with Image J (imagej.nih.gov/ij/) or MetaMorph (Molecular Devices, USA) analysis software. Quantitative analyses subtracted background from subject area fluorescence intensities, which can produce negative results in beads experiments in which background levels from latex are lower than those of mII oocytes.

### Statistical analysis

All experiments were performed on at least two days. The number of samples (*n*) per experiment reflects oocyte or embryo survival after manipulation. All samples were randomly collected; that is, we did not knowingly select different classes of healthy oocytes or embryos except where stated and no data were selectively excluded. Data analysis was performed with or without blinding. Statistical differences between pairs of data sets were analyzed by chi-squared or two-tailed unpaired *t*-test. Values of *p* < 0.05 were considered statistically significant.

## Acknowledgements

The authors gratefully acknowledge support to A.C.F.P. from the Medical Research Council, UK (MR/N000080/1 and MR/N020294/1), to A.C.F.P. and Y.-H.T. from the Biology and Biological Science Research Council, UK (BB/P009506/1) and to Y.-H.T. from the Wellcome Trust (200730/Z/16/Z). We thank Professor C. Tickle and Dr. M. VerMilyea for incisive comments during manuscript preparation, animal care laboratory staff at the University of Bath, and Professor M. Szczelkun for introducing the Tsai and Perry labs.

## Author contributions

A.C.F.P. and Y.-H.T. conceived core experiments, which were performed by T.S., M.A., S.G.P., L.Y.P.L and Y.-H.T. A.C.F.P. wrote the manuscript with comments from all authors.

## Additional information

**Supplementary Information** accompanies this paper.

## References

1. Mali, P., Yang, L., Esvelt, K. M., Aach, J., Guell, M., DiCarlo, J. E., Norville, J. E. & Church, G. M. RNA-guided human genome engineering via Cas9. Science 339, 823–826 (2013).

2. Cong, L., Ran, F. A., Cox, D., Lin, S., Barretto, R., Habib, N., Hsu, P. D., Wu, X., Jiang, W., Marraffini, L. A. & Zhang, F. Multiplex genome engineering using CRISPR/Cas systems. Science 339, 819–823 (2013).

3. Suzuki, T., Asami, M. & Perry, A. C. F. Asymmetric parental genome engineering by Cas9 during mouse meiotic exit. Sci. Rep. 4, 7621 (2014).

4. Esvelt, K. M., Smidler, A. L., Catteruccia, F. & Church, G. M. Concerning RNA-guided gene drives for the alteration of wild populations. eLife 3, e03401 (2014).

5. Pineda, M., Moghadam, F., Ebrahimkhani, M. R. & Kiani, S. Engineered CRISPR systems for next generation gene therapies. ACS Synth. Biol. 6, 1614–1626 (2017).

6. Dai, W. J., Zhu, L. Y., Yan, Z. Y., Xu, Y., Wang, Q. L. & Lu, X. J. CRISPR-Cas9 for in vivo Gene Therapy: Promise and Hurdles. Mol. Ther. Nucleic Acids 5, e349 (2016).

7. Burt, A. Site-specific selfish genes as tools for the control and genetic engineering of natural populations. Proc. R. Soc. Lond. B 270, 921–928 (2003).

8. Gantz, V. M. & Bier, E. Genome editing. The mutagenic chain reaction: a method for converting heterozygous to homozygous mutations. Science 348, 442–444 (2015).

9. Hammond, A., Galizi, R., Kyrou, K., Simoni, A., Siniscalchi, C., Katsanos, D., Gribble, M., Baker, D., Marois, E., Russell, S., Burt, A., Windbichler, N., Crisanti, A. & Nolan, T. A CRISPR-Cas9 gene drive system targeting female reproduction in the malaria mosquito vector Anopheles gambiae. Nat. Biotechnol. 34, 78–83 (2016).

10. Nihongaki, Y., Otabe, T. & Sato, M. Emerging approaches for spatiotemporal control of targeted genome with inducible CRISPR-Cas9. Anal. Chem. doi: 10.1021/acs.analchem.7b04757 (2016).

11. Hemphill, J., Borchardt, E. K., Brown, K., Asokan, A. & Deiters, A. Optical Control of CRISPR/Cas9 Gene Editing. J. Am. Chem. Soc. 137, 5642–5645 (2015).

12. Nihongaki, Y., Kawano, F., Nakajima, T. & Sato, M. Photoactivatable CRISPR-Cas9 for optogenetic genome editing. Nat. Biotechnol. 33, 755–760 (2015).

13. Richter, F., Fonfara, I., Bouazza, B., Schumacher, C. H., Bratovič, M., Charpentier, E. & Möglich, A. Engineering of temperature-and light-switchable Cas9 variants. Nucleic Acids Res. 44, 10003–10014 (2016).

14. Moreno-Mateos, M. A., Fernandez, J. P., Rouet, R., Vejnar, C. E., Lane, M. A., Mis, E., Khokha, M. K., Doudna, J. A. & Giraldez, A. J. CRISPR-Cpf1 mediates efficient homology-directed repair and temperature-controlled genome editing. Nat. Commun. 8, 20–24 (2017).

15. Davis, K. M., Pattanayak, V., Thompson, D. B., Zuris, J. A. & Liu, D. R. Small molecule-triggered Cas9 protein with improved genome-editing specificity. Nat. Chem. Biol. 11, 316–318 (2015).

16. Zetsche, B., Volz, S. E. & Zhang, F. A split-Cas9 architecture for inducible genome editing and transcription modulation. Nat. Biotechnol. 33, 139–142 (2015).

17. Nguyen, D. P., Miyaoka, Y., Gilbert, L. A., Mayerl, S. J., Lee, B. H., Weissman, J. S., Conklin, B. R. & Wells, J. A. Ligand-binding domains of nuclear receptors facilitate tight control of split CRISPR activity. Nat. Commun. 7, 12009 (2016).

18. Oakes, B. L., Nadler, D. C., Flamholz, A., Fellmann, C., Staahl, B. T., Doudna, J. A. & Savage, D. F. Profiling of engineering hotspots identifies an allosteric CRISPR-Cas9 switch. Nat. Biotechnol. 34, 646–651 (2016).

19. Liu, K. I., Ramli, M. N., Woo, C. W., Wang, Y., Zhao, T., Zhang, X., Yim, G. R., Chong, B. Y., Gowher, A., Chua, M. Z., Jung, J., Lee, J. H. & Tan, M. H. A chemical-inducible CRISPR-Cas9 system for rapid control of genome editing. Nat. Chem. Biol. 12, 980–987 (2016).

20. Cruz Silva, M. M., Madeira, V. M., Almeida, L.M. & Custódio, J. B. Hemolysis of human erythrocytes induced by tamoxifen is related to disruption of membrane structure. Biochim. Biophys. Acta 1464, 49–61 (2000).

21. Brauch, H., Mürdter, T. E., Eichelbaum, M. & Schwab, M. Pharmacogenomics of tamoxifen therapy. Clin. Chem. 55, 1770–1782 (2009).

22. Orias, F., Bony, S., Devaux, A., Durrieu, C., Aubrat, M., Hombert, T., Wigh, A. & Perrodin, Y. Tamoxifen ecotoxicity and resulting risks for aquatic ecosystems. Chemosphere 128, 79–84 (2015).

23. Chin, J.W. Expanding and Reprogramming the Genetic Code of Cells and Animals. Annu. Rev. Biochem. 83, 379–408 (2014).

24. Tsai, Y. H., Essig, S., James, J. J., Lang, K. & Chin, J. W. Selective rapid and optically switchable regulation of protein function in live mammalian cells. Nat. Chem. 7, 554–561 (2015).

25. Perry, A. C. F., Wakayama, T., Kishikawa, H., Kasai, T., Okabe, M., Toyoda, Y. & Yanagimachi, R. Mammalian transgenesis by intracytoplasmic sperm injection. Science 284, 1180–1183 (1999).

26. Perry, A. C. F., Rothman, A., de las Heras, J. I., Feinstein, P., Mombaerts, P., Cooke H. J. & Wakayama, T. Efficient metaphase II transgenesis with different transgene archetypes. Nat. Biotechnol. 19, 1071–1073 (2001).

27. Jinek, M., Chylinski, K., Fonfara, I., Hauer, M., Doudna, J. A. & Charpentier, E. A programmable dual-RNA-guided DNA endonuclease in adaptive bacterial immunity. Science 337, 816–821 (2012).

28. Li, X. Zhao, X., Fang, Y., Jiang, X., Duong, T., Fan, C., Huang, C. C. & Kain, S. R. Generation of destabilized green fluorescent protein as a transcription reporter. J. Biol. Chem. 273, 34970–34975 (1998).

29. Suzuki, T., Yoshida, N., Suzuki, E., Okuda, E. & Perry, A. C. F. Full-term mouse development by abolishing Zn^2+^-dependent metaphase II arrest without Ca^2+^ release. Development 137, 2659–2669 (2010).

30. Yoshida, N. & Perry, A. C. F. Piezo-actuated mouse intracytoplasmic sperm injection (ICSI). Nat. Protoc. 2, 296–304 (2007).

31. Suzuki, T., Asami, M., Hoffmann, M., Lu, X., Gužvić, M., Klein, C. A. & Perry, A. C. F. Mice produced by mitotic reprogramming of sperm injected into haploid parthenogenotes. Nat. Commun. 7, 12676 (2016).

32. Berta, P., Hawkins, J. R., Sinclair, A. H., Taylor, A., Griffiths, B.I., Goodfellow, P. N. & Fellous, M. Genetic evidence equating SRY and the testis-determining factor. Nature 348, 448–450 (1990).

33. Müller, G., Ruppert, S., Schmid, E. & Schutz, G. Functional analysis of alternatively spliced tyrosinase gene transcripts. EMBO J. 7, 2723–2730 (1988).

34. Sunagawa, G. A., Sumiyama, K., Ukai-Tadenuma, M., Perrin, D., Fujishima, H., Ukai, H., Nishimura, O., Shi, S., Ohno, R., Narumi, R., Shimizu, Y., Tone, D., Ode, K. L., Kuraku, S. & Ueda, H. R. Mammalian Reverse Genetics without Crossing Reveals Nr3a as a Short-Sleeper Gene. Cell Rep. 14, 662–677 (2016).

35. Philpott, C. C., Ringuette, M. J. & Dean, J. Oocyte-specific expression and developmental regulation of ZP3, the sperm receptor of the mouse zona pellucida. Dev. Biol. 121, 568–575 (1987).

36. Lewandoski, M., Wassarman, K. M. & Martin, G. R. Zp3-cre, a transgenic mouse line for the activation or inactivation of loxP-flanked target genes specifically in the female germ line. Curr. Biol. 7, 148–151 (1997).

37. Ernst, R. J., Krogager, T. P., Maywood, E. S., Zanchi, R., Beránek, V., Elliott, T. S., Barry, N. P., Hastings, M. H. & Chin, J. W. Genetic code expansion in the mouse brain. Nat. Chem. Biol. 12, 776–778 (2016).

38. Chen, Y. T., Ma, J., Lu, W., Tian, M., Thauvin, M., Yuan, C., Volovitch, M., Wang, Q., Holst, J., Liu, M., Vriz, S., Ye, S., Wang, L. & Li, D. Heritable expansion of the genetic code in mouse and zebrafish. Cell Res. 27, 294–297 (2017).

39. Han, S., Yang, A., Lee, S., Lee, H. W., Park, C. B. & Park, H. S. Expanding the genetic code of Mus musculus. Nat. Commun. 8, 14568 (2017).

40. Schaefer, K. A., Wu, W. H.,Colgan, D. F., Tsang, S. H., Bassuk, A. G. & Mahajan, V. B. Unexpected mutations after CRISPR-Cas9 editing in vivo. Nat. Methods 14, 547–548 (2017).

41. Konermann, S., Brigham, M. D., Trevino, A. E., Joung, J., Abudayyeh, O. O., Barcena, C., Hsu, P. D., Habib, N., Gootenberg, J. S., Nishimasu, H., Nureki, O. & Zhang, F. Genome-scale transcriptional activation by an engineered CRISPR-Cas9 complex. Nature 517, 583–588 (2015).

42. Vora, S., Tuttle, M., Cheng, J. & Church, G. Next stop for the CRISPR revolution: RNA-guided epigenetic regulators. FEBS J. 283, 3181–3193 (2016).

43. Liao, H. K., Hatanaka, F., Araoka, T., Reddy, P., Wu, M. Z., Sui, Y., Yamauchi, T., Sakurai, M., O'Keefe, D. D., Núñez-Delicado, E., Guillen, P., Campistol, J. M., Wu, C. J., Lu, L. F., Esteban, C. R. & Izpisua Belmonte JC. In Vivo Target Gene Activation via CRISPR/Cas9-Mediated Trans-epigenetic Modulation. Cell doi: 10.1016/j.cell.2017.10.025 (2017).

44. Liu, X. F. & Bagchi, M. K. Recruitment of distinct chromatin-modifying complexes by tamoxifen-complexed estrogen receptor at natural target gene promoters in vivo. J. Biol. Chem. 279, 15050–15058 (2004).

45. Sharma, D., Saxena, N. K., Davidson, N. E. & Vertino, P. M. Restoration of tamoxifen sensitivity in estrogen receptor-negative breast cancer cells: tamoxifen-bound reactivated ER recruits distinctive corepressor complexes. Cancer Res. 66, 6370–6378 (2006).

46. Rogers, C. S. Genetically engineered livestock for biomedical models. Transgenic Res. 25, 345–359 (2016).

47. Pawluk, A., Amrani, N., Zhang, Y., Garcia, B., Hidalgo-Reyes, Y., Lee, J., Edraki, A., Shah, M. Sontheimer, E. J., Maxwell, K. L. & Davidson, A. R. Naturally occurring off-switches for CRISPR-Cas9. Cell 167, 1829–1838.e9 (2016).

48. Charlesworth, C. T., Deshpande, P. S., Dever, D. P., Dejene, B., Gomez-Ospina, N., Pavel-Dinu, M., Camarena, J., Weinberg, K. I. & Porteus, M. H. Identification of Pre-Existing Adaptive Immunity to Cas9 Proteins in Humans. bioRxiv, doi: http://dx.doi.org/10.1101/243345 (2018).

49. Shao, Y., Guan, Y., Wang, L., Qiu, Z., Liu, M., Chen, Y., Wu, L., Li, Y., Ma, X., Liu, M. & Li, D. CRISPR/Cas-mediated genome editing in the rat via direct injection of one-cell embryos. Nat. Protoc. 9, 2493–2512 (2014).

